# Huntingtin Over-Expression Does Not Alter Overall Survival in Murine Cancer Models

**DOI:** 10.1101/2022.09.11.507440

**Authors:** Laura Chan, Austin Hill, Ge Lu, Jeremy Van Raamsdonk, Randy D. Gascoyne, Michael R. Hayden, Blair R. Leavitt

**Affiliations:** Department of Medical Genetics and Centre for Molecular Medicine and Therapeutics, Child and Family Research Institute, University of British Columbia, Vancouver, BC, Canada, V5Z 4H4; Department of Neurology and Neurosurgery, McGill University, Montreal, QC, Canada; Metabolic Disorders and Complications Program, and Brain Repair and Integrative Neuroscience Program, Research Institute of the McGill University Health Centre, Montreal, QC, Canada; Center for Lymphoid Cancer, British Columbia Cancer, Vancouver, British Columbia, Canada; Department of Pathology and Laboratory Medicine, University of British Columbia, Vancouver, British Columbia, Canada

**Keywords:** Huntington’s disease, Cancer, Tumorigenesis, Mouse models, Huntingtin, Apoptosis

## Abstract

A reduced incidence of various forms of cancer has been reported in Huntington’s disease (HD) patients and may be due to pro-apoptotic effects of mutant huntingtin. We tested this hypothesis by assessing the effects of huntingtin protein over-expression on survival in two murine cancer models. We generated yeast artificial chromosome (YAC) HD mice containing human huntingtin transgenes with various CAG tract lengths (YAC18, YAC72, YAC128) on either an *Msh2 or p53* null background, which have increased cancer incidence. In both mouse models of cancer, the over-expression of either mutant or wild-type huntingtin had no significant effect on overall survival. These results do not support the hypothesis that mutant huntingtin expression is protective against cancer.

## INTRODUCTION

Huntington’s disease (HD) is an autosomal dominant neurodegenerative disorder clinically characterized by progressive motor impairments, cognitive decline, and neuropsychiatric features [1,2]. HD is caused by CAG trinucleotide repeat mutations in the *HTT* gene that result in an expanded polyglutamine tract in the N-terminus of the huntingtin protein [3,4]. Huntingtin protein is critical to neurodevelopment and normally plays a role in a variety of cellular functions including transcriptional regulation, BDNF production, axonal transport, endosomal trafficking, and vesicular recycling [5]. Huntingtin may also have a pro-survival role in the adult CNS and the levels of wild-type huntingtin have been shown to influence the toxicity of mutant huntingtin [6,7].

Mutant huntingtin (mHTT) acts primarily through a toxic gain of function to disrupt a variety of cellular processes [8], it is expressed ubiquitously throughout the body [9], and causes neurodegeneration and neuronal loss through apoptosis [10]. Additionally, mHTT has been associated with apoptosis mechanisms caused by DNA instability [11], with DNA instability being a very common characteristic of many cancers [12]. It’s been hypothesized that mHTT expression might be protective against cancer by inducing or increasing the rate of apoptosis in preneoplastic cells, thereby halting or preventing the development or progression of malignant tumors [13].

Epidemiologic studies identified a decreased incidence of various forms of cancer in the HD population [14, 15, 16]. First described by Sorensen & Fenger (1992), a 5% rate of death from cancer in HD patients were recorded in comparison to the 31% rate of death in first degree relatives in a Danish population [14]. Ji et al., (2012) went on to report similar findings in a Swedish population, finding a decreased risk of cancer in HD and other polyglutamine (polyQ) diseases [15]. This association was replicated in a French population by a different study group in 2017 [16]. The expanded polyglutamine tract in mutant huntingtin may possibly be acting as a tumor suppressor, inducing apoptosis in cells where the genome is unstable, and potentially mirroring the mechanism of neuronal cell death in HD.

The proposed relationship between the mutation in HD and tumorigenesis may not be the case for all cancer types. Looking at common breast cancer mutations produced conflicting results dependent on the mutation type [17]. Certain cancers, like of the skin and digestive tract [14, 18], are not included in the decreased incidence, while prostate and colorectal cancers were calculated in the HD REGISTRY to have some of the lowest incidence rates [18]. The environment of signaling pathways and tissue context within different cancer subtypes appears to influence the interplay between mHTT and growth of cells. For example, a mouse study investigating polyomavirus middle T antigen (PyVT) oncogene with an HD mouse model, *Hdh*^*Q111/Q111*^, found that mice homozygous for the CAG expanded mouse huntingtin gene had accelerated disease in two breast cancer models and lung metastasis [19]. Utilizing a variety of mouse models will further our intersectional and individual knowledge of cancer and HD pathways.

Inactivation of the *Msh2* gene in mice results in a predisposition to cancer through mismatch repair deficiency [20]. Homozygous (*Msh2*-/-) mice develop lymphoid tumors from genetic instabilities starting at two months old, making it a reliable model to be study lymphoma [21]. While previous studies have examined the effect of Msh2 deletion on CAG somatic instability [22] and striatal degeneration [23] in HD mouse models, to our knowledge no previous studies of *Msh2* deficiency in HD models have examined survival outcomes from cancer.

p53 is another tumour suppressor gene that has causes cancer when deleted in mouse models. A p53 “knock-out” (*p53-/-)* mouse will develop many types of tumors by the average age of 4.5 months, commonly lymphoma. [24, 25]. N171-82Q HD transgenic mice on a p53 null background had decreased behavioral abnormalities [26] and another HD mouse model, *Hdh*^*Q140*^ mice, crossed with *p53-/-* mice exhibited decreased mHTT expression in some organs [27], and increased lifespan in those mice with the mHTT transgene [28]. This survival data from the *Hdh*^*Q140*^; *p53-/-* model has yet to be repeated using another HD mouse line.

Yeast artificial chromosome (YAC) transgenic HD mice express human *HTT* transgenes with different CAG repeat expansions that express huntingtin with differing polyglutamine lengths [29, 30]. YAC18 mice express human wild-type huntingtin in addition to mouse huntingtin and do not develop any HD phenotypes. In fact, these mice are protected from various forms of neuronal injury [7,31]. The CAG-expanded YAC72 line presents with selective striatal neurodegeneration similar to the neurodegeneration in HD [29]. Further CAG expansion in the YAC128 mice results in robust motor abnormalities, cognitive dysfunction and progressive neuropathology that resembles human HD [30, 32]. The diversity of CAG repeat lengths in the YAC *HTT* transgene of this well-established mouse model allows investigation into the effects of mutant and wild-type huntingtin expression on cancer survival.

This study investigated the role of huntingtin over-expression on cancer survival by crossing YAC HD mice with two different cancer models (*Msh2* and *p53* “knockout” mice). Specifically, we examined the lifespan of these mice lines using Kaplan-Meier survival curves. We had hypothesized that mutant huntingtin is protective in cancer by causing apoptosis and/or inhibiting the growth of cancer cells. However, here we demonstrate that mouse models containing the YAC HD transgene and deletions in two separate tumour suppressors genes did not differ in overall survival.

## MATERIALS AND METHODS

### Mouse crosses and survival

YAC transgenic mice (YAC18 line B60, YAC72 line 2511, YAC128 line 53), *Msh2-/-* and YAC HD; *p53-/-* mice, expressing wild-type (YAC18) and mutant (YAC 72, 128) human huntingtin and inactivation of tumour suppressing genes (Msh2, p53) were generated through a series of genetic crosses. (Figure 1A, B). Littermates that did not contain the YAC HD transgene were used as controls. Mouse strains were maintained on a pure FVB/N (Figure 1A) or F2 hybrid C57BL/6 (Figure 1B) background. Survival data on mice were obtained from our own colony. Life span of an animal is defined as the number of days between birth and when the animal requires humane sacrifice, becomes moribund, or dies of natural causes. The colony was maintained in a 12 h light/dark cycle with ambient temperature and humidity. All animal experiments were conducted in accordance with the ethical guidelines described in the *Guide for the Care and Use of Laboratory Animals by the National Research Council*, and review and approved by the Animal Care and Use Committee of the University of British Columbia.

**Figure 1.**
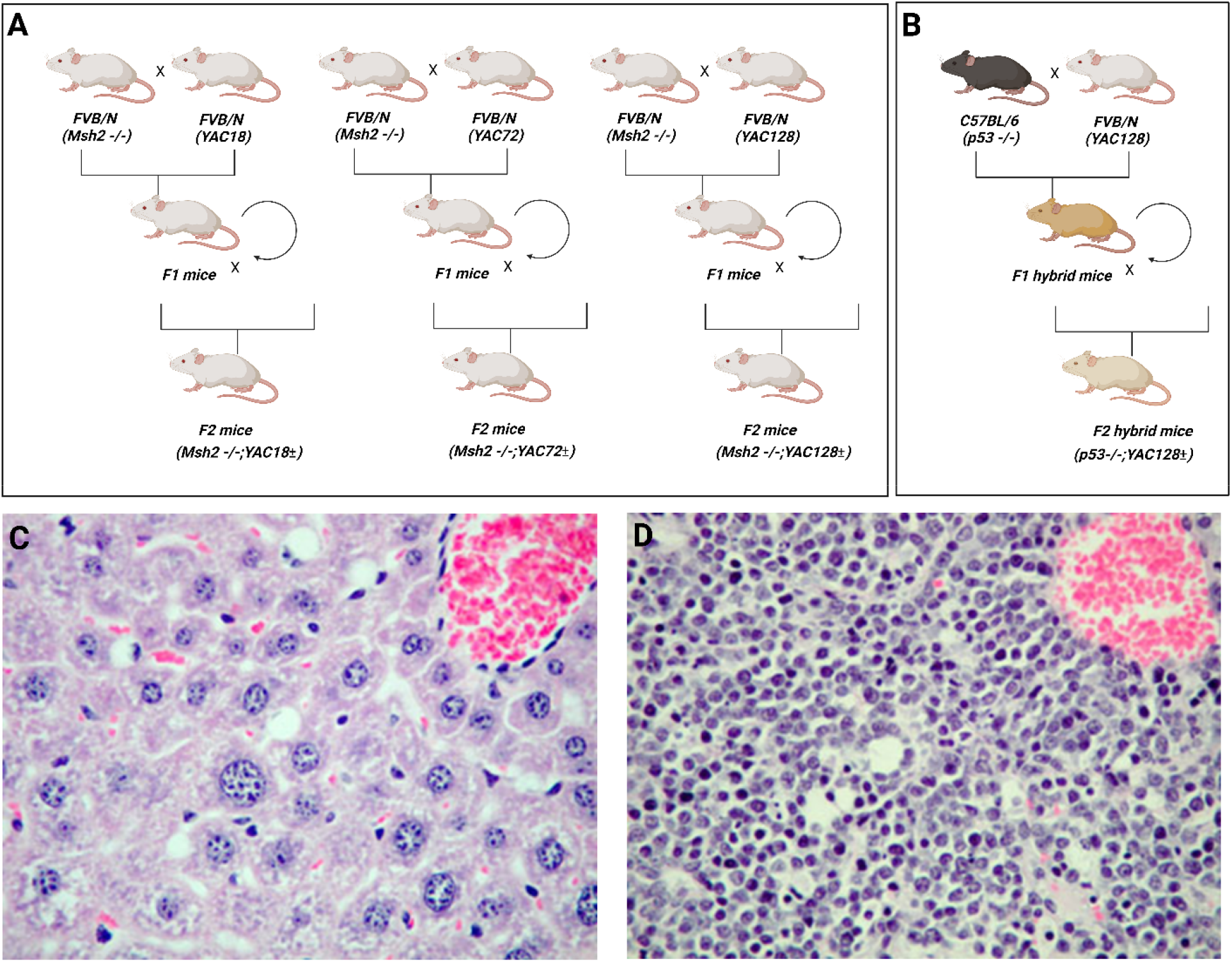
(A) Breeding scheme diagram for *Msh2-/-* mice with YAC HD mice (18, 72, 128) on a pure FVB/N background. (B) Breeding scheme diagram for *p53-/-* mice with YAC128 mice generating a F2 hybrid FVB/N x C57BL/6 background. (C) Normal liver histology of a wild-type *Msh2* +/+ mouse. (D) Liver histology of *Msh2*-/- mouse exhibiting lymphoblastic tumor consistent with lymphoma. Figure created with BioRender.com.

### Genotyping

Tail clippings were collected, genomic DNA prepared, and PCR amplification was carried out. The following primers were used to detect genotype: LYA1= 5’CCTGCTCGCTTCGCTACTTGGAGC 3′, LYA2= 5′ GTCTTGCGCCTTAAACCAACTTGG 3′, RYA1= 5′ CTTGAGATCGGGCGTTCGACTCGC 3′, RYA2=5′ CCGCACCTGTGGCGCCGGTGATGC 3′.

### Statistical analysis

Survival curves were generated then analyzed with a log rank, conducted in GraphPad Prism 9.0.1 for Windows, GraphPad Software, San Diego, California USA.

## RESULTS

To determine the effect of mutant and wild-type huntingtin on cancer survival, we crossed *Msh2-/-* mice to YAC transgenic mice (YAC18, YAC72, YAC128) and interbred them to obtain YAC HD+;*Msh2-/-* mice containing the human *HTT* gene with 18, 72, or 128 CAG repeats and YAC HD-;*Msh2-/-* controls (Figure 1A). Control mice lacked the human *HTT* transgene but were Msh2 deficient (YAC HD-;*Msh2-/-*). Mice with *Msh2* deficiency developed lymphoblastic tumors that infiltrated various organs such as the liver (Figure 1D). The gross pathology in the liver was confirmed to be consistent with lymphoma on histological sections by an experienced lymphoma pathologist.

Mice expressing a wild-type YAC *HTT* transgene (YAC18;*Msh2*-/-) mice had shorter median survival compared to control mice (Figure 1A). A log rank test performed on the survival curve of YAC18+;*Msh2-/-* (n=14) with control *Msh2-/-* littermates (n=10) mice were not statistically different (p=0.43) (Figure 2A). The median age of death was 129 days for mice containing the wildtype *HTT* transgene and 207 days for control mice with only the *Msh2-/-* mutation.

**Figure 2.**
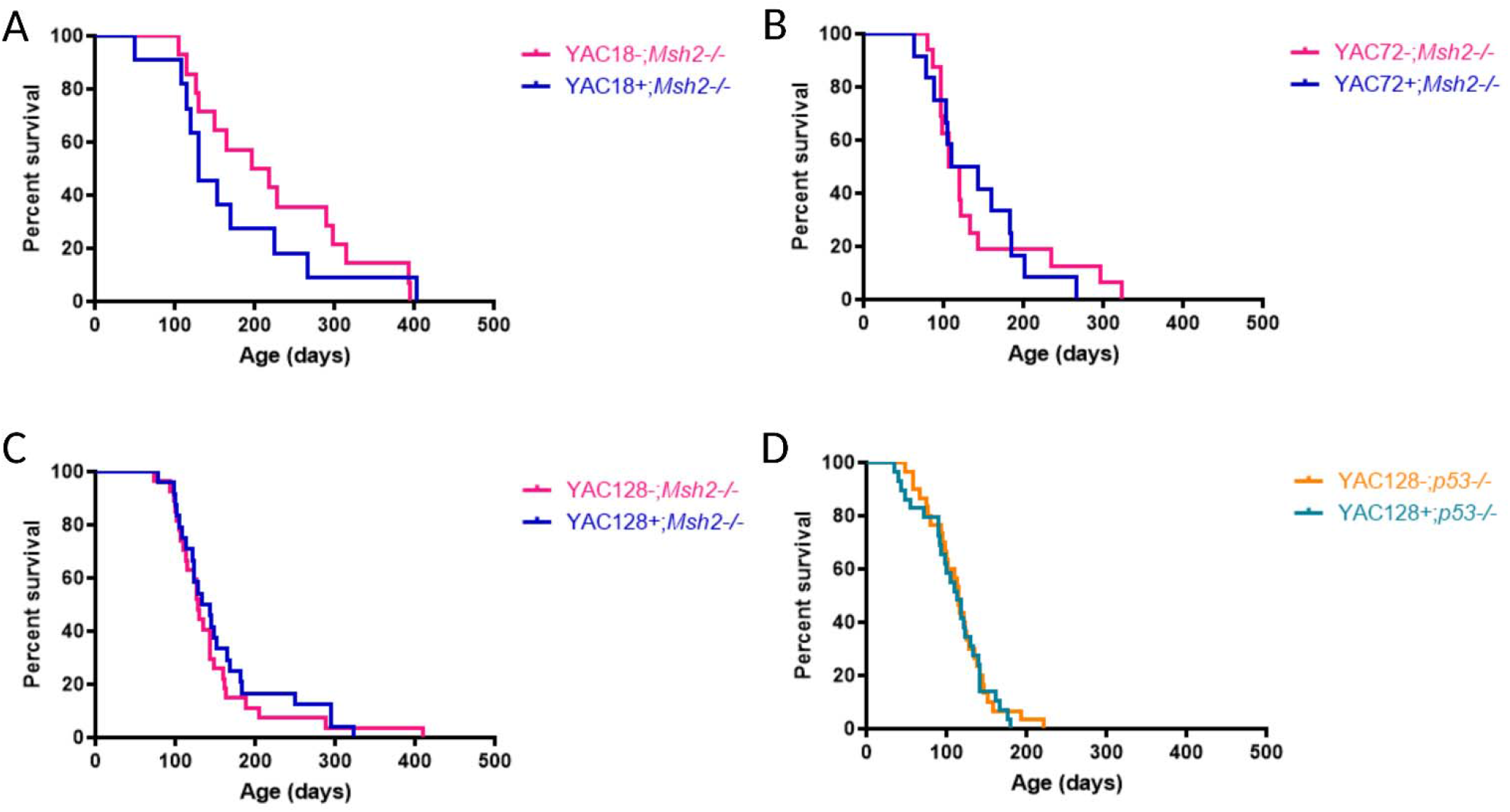
Kaplan-Meier survival curves of YAC HD mouse crosses using log rank test. Human HTT with varying CAG repeats expressing *Msh2* knockout mice, compared against control YAC HD-;*Msh2-/-* littermates: (A) YAC18; *Msh2*-/-. (B) YAC72; *Msh2-/-*, and (C) YAC128; *Msh2-/-*. (D) Human mHTT expressing *p53* knockout mice, compared against control YAC HD-;*p53-/-* littermates: YAC128; *p53-/-*. Graphs were generated using GraphPad Prism 9.0.1 for Windows, GraphPad Software, San Diego, California USA.

YAC HD mice expressing mutant huntingtin (YAC72 and YAC128) on the *Msh2-/-* background compared to YAC HD-;*Msh2-/-* mice also did not yield significant differences on survival in the following log rank tests. YAC72;*Msh2-/-* (n=11) had a median survival of 127 days compared to YAC HD-;*Msh2-/-* mice (n=16), with a slightly lower 113 days (p=0.92). (Figure 2B) Mice containing the YAC128 transgene with the longest CAG repeats YAC 128;*Msh2-/-* (n=23) had a similar result as the other YAC HD model, with a median survival of 138 days that was not statistically significant (p=0.49) from the YAC HD-;*Msh2-/-* mice (n=27) median survival of 127 days (Figure 2C). Survival curves between YAC HD+;*Msh2-/-* mice and control mice showed no significant differences in survival, which indicates that over-expression of mutant huntingtin does not alter survival in *Msh2-/-* mice.

In order to determine whether the lack of protection observed was specific to the *Msh2-/-* cancer model or whether a similar result would be obtained in other cancer models, we also crossed YAC128 mice to *p53-/-* cancer line mice (Figure 1B). Mice were genotyped and then overall survival assessed. A log rank comparison of survival curves of YAC128;*p53-/-* mice (n=30) with YAC HD-;*p53-/-* mice (n=29) found that there was not a statistically significant difference in survival (p=0.72) (Figure 2D). Median age of death for YAC128;*p53-/-* was 115 days, correspondingly the control mice were 112 days. This data indicates no survival benefit of mutant huntingtin expression in *p53-/-* mice.

## DISCUSSION

The overall survival of *Msh2-/-* and *p53-/-* mice was not significantly altered by the over-expression of mutant or wild-type huntingtin. There was no clear effect of different CAG repeat lengths for YAC18 mice expressing human wild-type huntingtin, nor for YAC72 and 128 mice expressing mutant huntingtin.

The potentially decreased median lifespan that was observed in *Msh2-/-* crossed with YAC18 mice compared to the Msh2-/- littermate controls (129 days versus 207), while not significant, is suggestive that increased wild-type huntingtin may accelerate oncogenesis and may help to explain some of the discrepant results in the literature [33]. Additional studies of the role of wild-type huntingtin in cancer are warranted.

There was no clear effect of mutant huntingtin over-expression on survival in these models. The interpretation of this mouse data is limited to the effects of over-expression of human mutant huntingtin, but implies that the presence of mHTT and CAG expansions do not have a significant impact on lifespan in mouse models of cancer, with the caveat that the results we have obtained may not apply to conditions in which the mutation gene occurs in the endogenous *HTT* gene. To our knowledge, this is the first study crossing different mHTT CAG repeat size mice models with tumour suppressor knockout mouse models. The results of our study appear to conflict with the survival benefit reported previously for the *Hdh* ^*Q140/Q140*^ HD mouse model on the *p53-/-* background with the median life span improving from 145 days to 171 days in mice homozygous for the large CAG expanded mouse alleles [28]. The over-expression of human mutant huntingtin from the YAC transgenes (∼128 maximum CAG length) in our models may not be sufficient to see an effect on lifespan in comparison to the 140 CAG length model used in that study.

Correlation between lymphoma and HD is not well established, and mutant huntingtin may not have impact apoptosis pathways in lymphoid tumorigenesis, possibly due to tissues specific factors [33]. Future research should assess the effects of huntingtin levels (both wild-type and mutant) on lifespan and tumorigenesis in different cancer models. Metastasis has been investigated in breast cancer with evidence of HTT expression influencing cancer progression [17,19,33]. Therefore, breast cancer mouse models may be good candidates for additional future studies. Investigating the shared apoptosis pathways between HD and cancer increases knowledge and comprehension of the molecular mechanisms by which both wild-type and mutant huntingtin may influence human health and disease.

## ACKNOWLEDGMENTS

We would like to thank all past members of the Leavitt Lab that contributed to this work. This research was supported by the University of British Columbia Faculty of Medicine Centre for Molecular Medicine and Therapeutics Transgenic facility and the Department of Medical Genetics.

